# Correlations between holistic processing, Autism quotient, extraversion, and experience and the own-gender bias in face recognition

**DOI:** 10.1101/492322

**Authors:** Mia Morgan, Peter J Hills

## Abstract

The variability in the own-gender bias (OGB) in face-recognition is thought to be based on experience (Herlitz & Lovén, 2013) and the engagement of expert face processing mechanisms for own-gender faces. Experience is also associated with personality characteristics such as extraversion and Autism, yet the effects of these variables on the own-gender bias has not been explored. We ran a face recognition study exploring the relationships between opposite-gender experience, holistic processing (measured using the face-inversion effect, composite face effect, and the parts-and-wholes test), personality characteristics (extraversion and Autism Quotient) and the OGB. Findings did not support a mediational account where experience increases holistic processing and this increases the OGB. Rather, there was a direct relationship between extraversion and Autism Quotient and the OGB. We interpret this as personality characteristics having an effect on the motivation to process own-gender faces more deeply than opposite-gender faces.

---

Faces provide essential information for social interactions, such as identity, mood, and sex (Kanwisher & Yovel, 2006). It is, therefore, not surprising that humans have developed expert processing abilities dedicated specifically to faces (Pascalis et al., 2011) and are able to accurately recognise thousands with ease (Hancock & Rhodes, 2008). However, this skill does not extend to all faces: individuals have better recognition for faces of their own ethnicity (Meissner & Brigham, 2001), age (Rhodes & Anastasi, 2011), and gender (Herlitz & Lovén, 2013) than the respective out-group. These own-group biases are a reliable effect in the face recognition literature, nevertheless theoretical explanations for these effects are not complete.

While a crossover interaction has been found in the own-ethnicity bias (Meissner & Brigham, 2001), the own-gender bias (OGB) is usually found to be stronger in females than males (Lewin & Herlitz, 2002; Lovén, Herlitz, & Rehnman, 2011; Rehnman & Herlitz, 2007). This is coupled with the results highlighting females have higher recognition accuracy for all faces than males (Lewin & Herlitz, 2002; Rehnman & Herlitz, 2007). This latter finding highlights an individual difference variable in face recognition. Such individual differences are not often considered in face recognition research, and in particular in the own-group biases. Potentially, while gender may affect face recognition, it is possible that a covarying individual difference variable may better account for the effect on face recognition. These have not been considered in detail in models of the own-group models.

The asymmetrical nature of the OGB makes establishing the underlying mechanisms potentially more difficult (Man & Hills, 2016) unless one considers individual variability. Many theoretical explanations for the OGB are based on theories explaining the own-ethnicity bias. These are usually based on the notion that extensive experience with own-group faces (which is one individual difference variable) results in heightened expert processing and this experience is lacking with out-group faces (Chiroro & Valentine, 1995) causing them to be less well recognised. Here we summarise the notion of expert processing and how it relates to the OGB and explore other individual difference variables that may play a role in the bias.

## Expert processing

Expert processing of faces is typically considered to be based on holistic processing (Maurer, Le Grand, & Mondloch, 2002): attending to the face as a Gestalt whole (Rossion, 2008) or processing the features in parallel (Palermi & Gauthier, 2012), developed from years of experience with upright faces. Michel, Corneille, and Rossion (2010) have indicated that own-ethnicity faces are processed with more holistic processing than other-ethnicity faces, indicating that the own-group biases are indeed based on expertise (see also De Heering & Rossion, 2008; Hills, Willis, & Pake, 2017; Michel, Rossion, Han, Chung, & Caldara, 2006; Michel, Caldara, & Rossion, 2006). Man and Hills (2016) measured whether more holistic processing was employed for own-gender faces than opposite-gender faces using the face-inversion effect (FIE: Yin, 1969). However, they found no difference in the FIE between own- and opposite-gender faces. These results indicate that there is no increased expertise afforded to own-gender faces relative to other-gender faces. However, the FIE is but one measure of holistic processing and may actually reflect the engagement of expertise rather than holistic processing per se.

While the FIE might measure the engagement of holistic processing, two further paradigms directly measure holistic processing (Maurer et al., 2002): The composite face effect (CFE: Young, Hellawell, & Hay, 1987); and the parts and whole test (Tanaka & Farah, 1993). In the CFE, participants perform worse at matching identical top halves of a face when they are paired with the bottom half of a different identity; however this effect disappears when the halves are misaligned (Young et al., 1987). This displays that face parts cannot be viewed in isolation and rather the bottom half of the face creates a whole new identity, demonstrating holistic processing (Rossion, 2013). In the parts and wholes test, parts of faces are identified more accurately when in a whole face, compared to in isolation (Tanaka & Farah, 1993). This whole-part advantage is considered a measure of holistic processing. Given that holistic processing is assumed to correlate with the own-group biases, we predict that holistic processing measured by these tasks would be greater for own-relative the opposite-gender faces.

## Experience dependent expert processing

We implied earlier that the expertise in face recognition is accrued through extensive experience. Given this, one would assume that cross-group experience would correlate with the magnitude of the own-group biases. This is the contact hypothesis (Bukach, Cottle, Ubiwa, & Miller, 2012; Chiroro & Valentine, 1995; Hancock & Rhodes, 2008; Harrison & Hole, 2009) and evidence for it has been found to explain the own-ethnicity bias (De Heering & Rossion, 2008; De Heering, De Liedekerke, Deboni, & Rossion, 2010; Michel et al., 2006a, b) and the own-age bias (Harrison & Hole, 2009; Kuefner, Macchi Cassia, Picozzi, & Bricolo, 2008). Furthermore, increased contact with faces of the out-group increases the amount of holistic processing afforded to those faces (Megias, Rzeszewska, Aguado, & Catena, 2018; Tanaka, Kiefer, & Buckach, 2004). If contact affects the amount of holistic processing engaged in, then we would anticipate that holistic processing use would mediate the relationship between experience and the OGB.

Evidence for the involvement of experience in the OGB is not convincing: Wolff, Kemter, Schweinberger, & Wiese, (2013) found no correlation between self reported daily contact with faces of the opposite gender and expert processing (see also Hills et al., 2018, who also measured objective experience due to schooling). While early interactions are usually female dominated (Culkin, 1999; Rennels & Davis, 2008) and this might lead to a bias, subsequent peer interactions in middle childhood (Rubin, Bukowski, & Parker, 1998) are usually same-sex (Bukowski, Brendgen, & Vitaro, 2007). Nevertheless, the role of contact in the OGB has not been conclusively precluded yet.

While contact explicitly might not be responsible for the OGB, it is entirely possible that gender differences in socialisation affect face-recognition performance more generally (Rehnman & Herlitz, 2007). Women have superior face recognition accuracy overall compared to men (Rehnman & Herlitz, 2007) due to increased socialisation (Oswald, Clark, & Kelly, 2004; Rehnman & Herlitz, 2006) and contact and this might lead to gender differences in the deployment of holistic processing (Collishaw & Hole, 2000). Such an argument suggests anything that is associated with increased socialisation, such as extraversion, will lead to higher face recognition performance. Furthermore, variables that might correlate with the amount of holistic processing will correlate with the magnitude of the own-group bias. One such individual difference variable to consider is Autism Quotient.

Extraversion is characterised by interest in interpersonal relationships, social dominance and assertiveness (Depue & Collins, 1999). Given its association with socialisation and the theory posited above, we would assume that there will be a correlation between extraversion and face recognition accuracy. Indeed, Li et al. (2010) have found that extraverts show better face recognition than introverts based on the idea that they are more interested in faces than introverts. The theoretical argument for why women show enhanced face recognition relative to men is based on the same socialisation mechanisms. Therefore, we can suggest that extraverts will show enhanced face recognition relative to introverts and this is likely to be based on holistic processing given its involvement in expert face recognition, leading to a larger OGB.

Autism is a neurodevelopmental disorder associated with impairments in socialising and communication (Miles et al., 2005). Core traits of autism include a lack of socialisation (LaGasse, 2017), decreased reliance on global processing which is akin to holistic processing (Behrmann, Thomas & Humphreys, 2006), and higher levels of introversion (Oxford & Lavine, 1991). Given the thesis so far, these features would lead to lower face recognition performance. Indeed, Weigelt, Koldewyn, and Kanwisher (2012) have shown that individuals with autism have poorer face recognition accuracy than those without. This impairment has been attributed to reduced reliance on holistic processing (Joseph & Tanaka, 2003; Teunisse & de Gelder, 2003) which maybe a consequence of reduced exposure to faces throughout development. Evidence displays that males show higher autism traits than females (Baron-Cohen, Wheelwright, Skinner, Martin, & Clubley, 2001), which may potentially explain men's inferiority in face recognition relative to women. Further, the lack of socialisation would therefore lead to the hypothesis that autism would negatively correlate with the OGB.

## The present study

We have presented an argument that the OGB in face recognition is based on increased experience with faces of one's own gender relative to the opposite gender. This experience leads to differential amounts of holistic processing being engaged in for faces of one's own-gender and faces of the opposite gender. In this study, we will measure the effect of experience on the OGB, while assuming holistic processing acts as a mediator in this relationship. We will directly measure holistic processing using the CFE (Tanaka & Farah, 1993) and the parts and whole test (Young et al., 1987) as well as the FIE (Yin, 1969) for both own- and opposite-gender faces. We anticipate the amount of experience with faces of the opposite gender (specifically during infancy) will correlate with face recognition accuracy and that this relationship will be mediated by the level of holistic processing engaged in. Based on the same logic, we assume that there will be a correlation between extraversion and Autism Quotient and the OGB mediated by holistic processing and experience in a neurotypical sample.

## Method

### Participants

An opportunity sample of 82^1^ participants took part in the study: 40 males (aged 18 to 35 years, mean: 21), and 42 females, (18 to 47 years, mean: 20). Sample size was determined based on the effect size of the own-gender bias (Herlitz & Lovén, 2011).

With an *r*=.27, assuming a power of .8, sample size required was 81, as calculated using G-Power version 3.1. Participants took part as a partial fulfilment of a course requirement.

### Design

A correlational design with mediation was used. Across hypotheses, there were three predictor variables: experience with the opposite gender measured with the social contact questionnaire (Walker & Hewstone, 2006), extraversion measured using the Eysenck Personality Questionnaire Revised - short scale (Eysenck, Eysenck, & Barrett, 1985) and autism measured with the Adult Autism Spectrum Quotient (AASQ: Baron-Cohen et al., 2001). The mediator variable was holistic processing, measured using the CFE (Young, Hellawell, & Hay, 1987), the parts and whole test (Tanaka & Farah, 1993), and the FIE (Yin, 1969). The dependent variable was the magnitude of OGB (calculated using Hills & Lewis’, 2006 method: own gender *ď*-other gender *ď*)/(own gender *ď*-other gender *ď*), using accuracy measured in terms of Signal Detection Theory (*ď*; Green & Swets, 1966 calculated using the MacMillan and Creelman, 2005, method).

### Materials

#### Faces

For all tasks, the faces used were of White ethnicity, presented in frontal poses, with neutral or smiling expressions. All images were grey-scaled, cropped to remove neck and hair length, then pasted onto a white background. For the recognition task, two versions of 80 face images (40 male and 40 female; aged between 18 and 30 years) were taken from the Minear and Park (2004) database: One was displayed at learning and one was displayed at test (this was counterbalanced across participants). Inverted faces were created by rotating the stimuli 180 degrees. Images were resized to be 1024 px by 768 px presented in 300 dpi resolution. This was all performed using GNU Image Manipulation Program (GIMP; version 2.8.10). The images used at learning and test were counterbalanced between participants.

For the composite faces, a further 12 (6 female) faces were taken from the Minear and Park (2004) database. Instead of combining parts of different faces, like the standard task (Young et al., 1987), different identities were created by moving features to increase the chance of participants using holistic processing; as position was the only aspect of the features that changed (Lewis & Hills, 2018). Twelve new images were created by adjusting the distance between the eyes, and a further 12 by moving the mouth up. A further 12 images included both the changes. An additional 48 images were created by vertically splitting the image through the nose and moving the lower half of the face to the right (see Lewis & Hills, 2018, Figure 3). This resulted in 96 images.

For the Parts and Wholes test (Tanaka & Farah, 1991), stimuli consisted of 80 original faces (40 female) taken from the Minear & Park (2001) database. These images were cropped to create part images. Parts were created for the eyes, nose, and mouth (however, for the purpose of the experimentation, only trials involving the eyes were analysed because the whole-part advantage is much stronger for the eyes than other features, Joseph & Tanaka, 2002). Other features were included to prevent participants only focusing on the eyes (Michel et al., 2006).

#### Social contact questionnaire

This was adapted from Walker and Hewstone’s (2006) scale that measured experience with the own-ethnicity. Questions associated with ethnicity were modified to ask about experience with both genders. It consisted of 13 items for each gender measuring contact with people of each gender during infancy, childhood, and adulthood. Participants respond on a 5-point Likert scale, with anchor points ranging from strongly agree to strongly disagree or to indicate a numerical value. This produced an experience score for interactions with both males and females. Higher scores represented more social contact. The original scale had high internal reliability (Cronbach’s α= 0.83), construct, and face validity (Walker & Hewstone, 2006).

#### Eysenck Personality Questionnaire Revised

The short scale version of Eysenck’s Personality Questionnaire-Revised (EPQ-R; Eysenck et al., 1985) was used in this study. It consisted of 48 items measuring extraversion/introversion, neuroticism, psychoticism, and lying. Participants respond with a “yes” or a “no” to each statement. Only the 12 extraversion items were analysed, with higher scores representing more extraversion. This scale has high internal reliability (Cronbach’s α=.89-.91; Weaver & Kiewitz, 2007) and factorial validity (Wilson & Doolabh, 1991).

#### The Adult Autism Spectrum Quotient

The Adult Autism Spectrum Quotient (AASQ; Baron-Cohen et al., 2001) consisted of 50 items measuring traits of Autism. Participants responded using a 4-point Likert scale with anchor points ranging from “definitely agree” to “definitely disagree". This has moderate to high internal reliability (Cronbach’s α= 0.63-0.77) high test-retest reliability and criterion validity (Baron-Cohen et al., 2001). This created a score out of 50, with higher numbers representing more traits associated with Autism.

### Procedure

After providing informed consent, participants completed the three face recognition tasks (with embedded three questionnaires). Participants were offered breaks between each task. The order of tasks were counterbalanced between participants and once completed, participants were debriefed. The face tasks were displayed on a 15” high-resolution colour LCD.

#### Face-recognition task/Inversion effect

This task was completed on OpenSesame (Mathôt, Schreij, & Theeuwes, 2012) consisted of three consecutive phases: Learning, distraction, and test. At learning, participants were presented with 40 faces on screen, (20 female, 10 of which were inverted) sequentially in a random order in the centre of the screen. The face remained on screen until responded to by a distinctiveness judgement. Distinctiveness was measured by participants responding to the question, “How easy would this face be to spot in a crowd?” (Light, Kayra-Stuart, & Hollander, 1979), using a Likert scale ranging from 1 (difficult - i.e., typical face) to 9 (easy - i.e., atypical) This judgement was carried out as it is a gender-neutral form of encoding (Hills et al., 2018). Between each face a grey mask was presented in the centre of the screen where the face would appear for 600 ms.

In the distractor phase, participants completed the social contact questionnaire (Walker & Hewstone, 2006), EPQ-R (Eysenck et al., 1985), and AASQ (Baron-Cohen et al., 2001) on screen, in addition to providing demographic information. This typically lasted twenty minutes to complete.

At test participants were presented with 80 images of faces: 40 previously seen and 40 new images (split evenly by gender and orientation). Orientation of the old faces was matched from learning to test. Faces were presented sequentially in a random order. Between each face, there was an inter-trial stimulus consisting of a grey box presented in the centre of the screen where the face would appear for 1150 ms. The participants had to judge whether they recognised the face from the learning phase, pressing the ‘m’ computer key if they did and ‘z’ if not. Faces were on screen until participants responded.

#### Composite face effect

This task was completed using OpenSesame. This was a delayed-matching task. Participants were presented with 384 trials sequentially. Each trial consisted of a face presented for 600 ms, a blank screen for 300ms, followed by a second face presented until participants responded to it: participants had to judge whether the distance between the eyes in the first image were the same as the second image - pressing ‘m’ for similar and ‘c’ for different. This is akin to identifying if the two images represent the same identity.

Based on the pairing of the images (different eye and moth positions), there were 16 different types of pairing for both the aligned and the misaligned conditions. Therefore, there were 32 different trial types created for each of 12 face identities, creating a total of 384 trials. The order of the pairs were randomised between participants. This pairing style created four conditions conceptually for the aligned and misaligned variants: congruent same (CS), congruent different (CD), incongruent different (ID), incongruent same (IS). In two of the conditions, the bottom half of the face changed, and either the top half also changed (CD) or stayed the same (IS). In the other two conditions, the bottom half of the face stayed the same and the top half either changed (ID) or remained the same (CS). The metrics for calculating holistic processing are presented in Table 1.

#### Parts and whole test

This was a delayed-matching task. Participants were presented with 480 trials sequentially. Each trial consisted of a face for 1508ms, a blank inter-stimulus interval for 305ms, followed by a pair of images including the original image and a distractor face presented until participants responded to indicate with right and left computer keys which image they had previously seen. There was a blank inter-trial interval of 796ms. There were four blocks of 120 trials. Each block contained a different combination of images (from learning to test): whole-whole(WW), whole-part (WP), part-whole (PW), part-part (PP). Having four conditions, instead of the original two used by Tanaka and Farah (1993) (WW and WP) was used to give a more complete measure of holistic processing as it allows for relative performance differences between the part and whole conditions not to affect the calculation of holistic processing (Leder & Carbon, 2005). The metric for calculating holistic processing is presented in Table 1.The presentation order of the blocks and the images within the blocks were randomised between participants.

## Results

Before assessing our hypotheses, we first established that we had an OGB using a one-sample *t*-test, *t*(159)=3.56, *p*<.001. The OGB was larger for female participants than male participants, *t*(158)=1.98, *p*=.049. We then assessed whether each test of holistic processing produced results as expected using a series of one-sample *t*-tests for both own- and opposite-gender faces. This is presented in Table 1. We assessed whether these measures of holistic correlated with each other using a series of Bonferroni-corrected (α=.008) Pearson’s correlations. There were no significant correlations post-correction (largest *r*=.16, smallest *p*=.046)^2^, replicating Rezlescu, Susilo, Wilmer, and Caramazza (2017). In order to assess whether participants showed greater holistic processing for own-gender versus opposite-gender faces, we ran a series of within-subjects t-tests, also presented in Table 1. These all show enhanced holistic processing for own-gender than opposite-gender faces, except when measured using the parts and wholes test.

In order to address our main hypotheses (that experience would predict recognition through holistic processing), we ran a series of mediation analyses using the Baron and Kenny (1986) method. This analysis contains three linear regressions: regressing the predictor on the mediator; the predictor on the outcome variable; and both the predictor and mediator on the outcome variable, in a single linear regression. Given that our four measures of holistic processing did not correlate consistently with each other, we ran four mediations, with each holistic processing measure included. If all regressions displayed that the predictor significantly affects the outcome variable, Sobel’s test of mediation (Sobel, 1982) will be carried out.

Since experience (measured as a relative measure of experience with one’s own-gender and the opposite gender) did not predict the OGB, *R*(158)=.09, *F*=1.40, *MSE*=0.40, *p*=.238, we did not run the mediation analysis for this variable. We ran the analysis on both females and males separately as well, and neither reached significance, largest *R*(78)=.11, smallest *p*=.329. Nevertheless, we did assess whether measures of holistic processing predicted the OGB. We ran a series of Bonferroni-corrected (α=.0125) regressions between each measure of holistic processing and the magnitude of the OGB. None of these reached significance, largest *R*(158)=.15, smallest *p*=.066.

These measures were conducted only on the overall measure of experience. Since Hills et al (2018) have shown that recent experience with faces does not predict the OGB, it is no surprise that we failed to find a relationship between experience and the OGB. Therefore, we recalculated the experience questionnaire including only measures of childhood experience to assess whether self-reported childhood experience predicted the OGB.

**Table 1.** One sample t-test values, mean difference, and significance value, for each holistic processing measure for own- and other-gender faces. Within-subjects t-test values, mean difference, and significance for the OGB in these measures of holistic processing for female and male participants separately and overall.

| Test and authors | Test Metric | Face Gender | Mean difference | t value | Significance value (p) |
| --- | --- | --- | --- | --- | --- |
| Face-Inversion Effect (Yin, 1969) | (d' of upright faces)- (d' of inverted faces) | Own | 0.84 | 10.97 | .001 |
|  |  | Opposite | 0.55 | 7.13 | .001 |
|  |  | OGB<sub>female</sub> | 0.39 | 2.65 | .010 |
|  |  | OGB<sub>male</sub> | 0.20 | 1.35 | .180 |
|  |  | OGB | 0.29 | 2.84 | .005 |
| Partial composite effect (Rossion, 2013) | (mean: IS + CD aligned) – (mean: IS + CD misaligned) | Own | 0.59 | 22.38 | .001 |
|  |  | Opposite | 0.17 | 10.78 | .001 |
|  |  | OGB<sub>female</sub> | 0.36 | 7.74 | .001 |
|  |  | OGB<sub>male</sub> | 0.49 | 13.16 | .001 |
|  |  | OGB | 0.42 | 14.11 | .001 |
| Complete composite effect (Richler, Cheung & Gauthier, 2011)<sup>3</sup> | (mean: CS+CD aligned)-(mean: IS+ID aligned) | Own | 0.58 | 20.11 | .001 |
|  |  | Opposite | 0.15 | 8.45 | .001 |
|  |  | OGB<sub>female</sub> | 0.40 | 8.87 | .001 |
|  |  | OGB<sub>male</sub> | 0.46 | 10.44 | .001 |
|  |  | OGB | 0.43 | 13.64 | .001 |
| Parts and whole effect (Tanaka & Farah, 1993) | (WW-WP, Tanaka Farah, 1993) + (PP-PW, Leder & Carbon, 2005) | Own | 0.07 | 6.71 | .001 |
|  |  | Opposite | 0.09 | 8.50 | .001 |
|  |  | OGB<sub>female</sub> | -0.05 | 3.03 | .003 |
|  |  | OGB<sub>male</sub> | 0.01 | 0.50 | .618 |
|  |  | OGB | -0.02 | 1.74 | .084 |

<sup>3</sup> Two values were produced from the CFE, as both partial (Rossion, 2013) and complete analysis (Richler, Cheung & Gauthier, 2011) were run because of the ongoing debate as to which calculation represents holistic processing (Rossion, 2013) and studies have found holistic processing in one but not the other (Konar, Bennett & Sekuler, 2010; Richler et al., 2011).

Early experience did not predict the OGB, *R*(158)=.11, *F*=2.05, *MSE*=0.40, *p*=.154. Therefore, we did not run further mediation analyses for this variable. We ran the analysis on both females and males separately, and neither reached significance, largest *R*(78)=.11, smallest *p*=.342. Nevertheless, we did assess whether measures of holistic processing predicted the OGB.

Given that we have not found that experience is related to the OGB, we turned to look at the other individual difference variables we measured might be related to it. Turning first to Autism, we found that Autism predicted the OGB, *R*(158)=.33, *F*=19.03, *MSE*=0.36, *p*<.001. Social experience did not predict Autism, *R*(158)=.06, *F*=0.50, *MSE*=35.47, *p*=.495. Autism did predict holistic processing as measured by the FIE, *R*(158)=.31, *F*=16.52, *MSE*=0.47, *p*<.001 (but no other measures of holistic processing, largest *R*(158)=.07, smallest *p*=.382). The relationship between the FIE and the OGB was not significant as described above, *R*(158)=.15, *F*=3.42, *MSE*=0.39, *p*=.066, nevertheless, we ran the fourth regression with both the predictor (Autism) and the mediator (FIE) to assess any potential mediation. In this analysis, the effect of Autism on the OGB remained significant, *t*(158)=3.95, *p*<.001, indicating no significant mediation. Indeed, the Sobel test was not significant, Sobel=0.62, *p*=.53.

Using a parallel analysis protocol, we found that Extraversion predicted the OGB, *R*(158)=.36, *F*=23.52, *MSE*=0.35, *p*<.001. Social experience did not predict Extraversion, *R*(158)=.06, *F*=0.54, *MSE*=13.07, *p*=.465. There were non-significant trends for extraversion to predict holistic processing as measured by the FIE, *R*(158)=.13, *F*=2.87, *MSE*=0.51, *p*=.092, and the partial CFE, *R*(158)=.14, *F*=3.18, *MSE*=0.24, *p*=.077 (but not the other measures of holistic processing, largest *R*(158)=.02, smallest *p*=.817). As before, we ran the fourth regression with both the predictor (extraversion) and the mediators (FIE and partial CFE done separately) to assess any potential mediation. In these analyses, the effect of extraversion on the OGB remained significant when controlling for holistic processing measured by the FIE, *t*(158)=4.64, *p*<.001, and by the partial CFE, *t*(158)=4.73, *p*<.001. Sobel tests revealed no significant mediations, Sobel=1.03, *p*=.30, for the FIE mediation, and Sobel=0.44, *p*=.66, for the partial CFE mediation. The full dataset is available at http://bordar.bournemouth.ac.uk/20/.

## Discussion

The first aim of this study was to investigate whether experience with own-gender faces, specifically childhood and peer relationships, predicted the OGB. No relationship was found, consistent with past research (Hills et al., 2018; Wolff et al., 2013). Such replicated null results suggest that the contact hypothesis (Chiroro & Valentine, 1995), even when early relationships are considered, cannot account for the OGB. The contact hypothesis has successfully explained other biases (own-ethnicity, De Heering & Rossion, 2008 and own-age, Harrison & Hole, 2009). The reason why experience does not explain the OGB may result from the nature of the relationships during development. There is a switch from female dominated childhood relationships to own gender peer relationships. This switch might negate the effects of experience at least in boys (Bukowski et al., 2007; Rennels & Davis, 2008). Further, unlike the other biases, there is a much more equivalent contact with own- and other-gender faces: any effect of contact is mitigated due to a smaller relative difference in contact between own- and other-gender faces relative to the other biases.

We hypothesised that greater holistic processing would be observed for own- than other-gender faces and that this would mediate the relationship between experience and the OGB. The majority of measures revealed enhanced holistic processing for own compared to opposite-gender faces. This indicates that we automatically categorise faces as in-group or out-group (own- or other-gender in this case). In-group faces are then processed deeply, (i.e., holistically). Out-groups are attended to less resulting in shallower processing, as explained in the socio-cognitive motivational account (Sporer, 2001). Motivation can manipulate whether expert processing is applied (Hills et al, in press), so it maybe that the OGB is a result of motivation to process faces differently as opposed to experience. However, enhanced holistic processing for own-gender faces we found that it did not predict the OGB. This is consistent with findings from Konar, Bennett & Sekuler (2010) who found that holistic processing was not correlated with face recognition accuracy.

For female participants, increased holistic processing was found for own-gender faces relative to other-gender faces. This was not found across all measures for male participants. This might be related to the oft found result that the OGB is stronger in females than males (Lewin & Herlitz, 2002; Lovén et al., 2011; Rehnman & Herlitz, 2007). We link this finding to the notion that females are typically more interested in faces than males (Rehnman & Herlitz, 2006) which causes them to be more accurate at face recognition generally and show an enhanced OGB and holistic processing than males. Therefore, while we have not found that experience leads to holistic processing and this leads to the OGB, we have suggested that the mechanisms behind increased holistic processing and face recognition accuracy in females relative to males is due to the same mechanism.

One theoretical reason why females are thought to be better at face recognition than males is because they are socialised to be. In the introduction, we indicated that there might be other individual difference variables that are related to sociability that might explain the OGB. We predicted that there would be individual differences variables (extraversion and autism) affecting the amount of holistic processing engaged in and this would affect face recognition. We did not find this indirect mediational link between these variables. However, we found a direct relationship between autism and the OGB and between extraversion and the OGB. We did not predict the direct link between these individual difference variables and the OGB. Our results indicate we can discount the mediational link and explore alternative explanations for why these individual difference variables might affect face recognition.

Autism is known to affect perception and information processing profoundly (e.g., Dakin & Frith, 2005), including enhanced visual target detection (Berton, Mottron, Jelenic & Faubert, 2005) and abnormal eye movements when viewing social scenes (Fletcher-Watson, Leekam, Benson, Frank, & Findlay, 2009) including faces (Pelphrey, Sasson, Reznick, Paul, Goldman, & Piven, 2002). Given that Autistic traits exist along a continuum in the general population (Robinson et al., 2011), it is reasonable to consider that these effects also exist as a continuum in the general population. People with Autism are generally less accurate at face recognition than people without Autism (Weigelt et al., 2012). Riby and Hancock (2009) hypothesise that this is due to a decreased interest in socialisation associated with Autism. Our correlation between Autism Quotient and the OGB reveals even lower face recognition performance for opposite-gender faces compared to own-gender faces. This suggests that Autism levels correlate with the amount of attention paid to other-gender faces negatively. We hypothesise that this is related to the findings that individuals with Autism find it easier to communicate and interact with in-group than out-group members (Gernsbacher, Stevenson, & Dern, 2017). It is therefore reasonable to suggest that people with Autism pay less attention to out-group members (in this case opposite-gender faces) which leads to some level of cognitive disregard and therefore shallow processing.

While extraversion is primarily associated with an increase in sociability (Caspi, Roberts, Shiner, 2005), it is also known to affect perceptual processes, for example increased olfactory sensitivity (Koelega, 1970) and eye movements (Rauthmann, Seubert, Sachse, & Furtner, 2012) when viewing social scenes (Moss, Baddeley, & Canagarajah, 2012). Indeed, the effects extraversion has on eye movements may explain female accuracies in classification tasks (Moss et al., 2012). Extraversion correlates positively with face recognition accuracy (Li, et al., 2010). However, it does not correlate with recognition accuracy for non-social stimuli (Li et al., 2010). Our data demonstrated that extraversion was associated with poorer recognition of other-gender faces (because of the larger OGB). This was not related to experience (potentially therefore sociability) nor increased use of holistic processing. Instead, we hypothesise that this relationship occurred due to extraverts increased attention paid to faces relative to introverts as revealed through the ERP P300 (Fishman, Ng, & Bellugi, 2011). This ERP is known to be stronger for self-relevant stimuli than non-self relevant stimuli (Gray, Ambady, Lowenthal, & Deldin, 2004; Ninomiya, Onitsuka, Chen, Sato & Tashiro, 1998). Therefore, we suggest that extraverts pay more attention to their own-gender faces because they are more willing to engage in socialisation with their own-group (e.g., through sporting or social activities, Paunonen, 2003) and this makes their own-group more important to process.

These results highlight a hitherto under-researched area of face recognition research, that of individual differences associated with personality traits. While it is known that extraversion and Autistic traits affect face recognition generally (Gelder, Vroomen, & Van der Heide, 1991; Li et al., 2010), what we have shown is that individual differences can have more nuanced effects on face recognition. These individual difference variables appear to be dimensional in their effect on face recognition given the correlational nature of the present study. Furthermore, individual difference variables might have different effects on different classes of faces. These results are important in building models of face recognition and might aid researchers when theorising how face recognition develops.

In conclusion, we have replicated the OGB in face recognition, demonstrating that it is stronger in males than females. Females also consistently show more holistic processing for own-gender faces than other-gender faces, however this is unrelated to both experience with own- and opposite-gender faces and to the OGB. Further, we have shown that the magnitude of the OGB is predicted by participants’ Autism and extraversion scores. These results show subtle individual difference variability affects reliable face recognition findings.

## Footnotes

1 Originally, there were five additional participants whose data were excluded because they did not complete the task appropriate - by pressing a single key throughout the experiment

2 Pre-correction, the complete CFE correlated negatively with the whole-part advantage, *r*(156)=.16, *p*=.046 and a negative correlation between the FIE and the partial CFE, *r*(158)=.14, *p*=.046. No other correlations were significant, largest *r*=.04, smallest *p*=.348.

